# How evolution draws trade-offs

**DOI:** 10.1101/169904

**Authors:** Salomé Bourg, Laurent Jacob, Frédéric Menu, Etienne Rajon

## Abstract

Recent empirical evidence suggest that trade-off shapes can evolve, challenging the classical image of their high entrenchment. Here we model the evolution of the physiological mechanism that controls the allocation of a resource to two traits, by mutating the expression and the conformation of its constitutive hormones and receptors. We show that trade-off shapes do indeed evolve in this model through the combined action of genetic drift and selection, such that their evolutionarily expected curvature and length depend on context. In particular, a trade-off’s shape should depend on the cost associated with the resource storage, itself depending on the traded resource and on the ecological context. Despite this convergence at the phenotypic level, we show that a variety of physiological mechanisms may evolve in similar simulations, suggesting redundancy at the genetic level. This model should provide a useful framework to interpret and link the overly complex observations of evolutionary endocrinology and evolutionary ecology.

## INTRODUCTION

Evolutionary biologists have long known that heritable characters (or traits) usually do not vary independently from each others (Stearns, 1992; Roff, 1993). Such pleiotropy is thought to hinder adaptive evolution, both because it restricts the available combinations of traits and because mutations are less likely favorable when they impact many traits (Fisher, 1930; Orr, 2005; Paaby & Rockman, 2013; Hine *et al.*, 2014). How pleiotropy evolves, and whether it is universal – *i.e.* every gene contribute, to some extent, to all traits – or restricted is the subject of an old but still ongoing debate (Wright, 1980; Wagner *et al.*, 2008; Wagner & Zhang, 2011; Hill & Zhang, 2012; Wagner & Zhang, 2012).

For many traits – when scaled such that they correlate positively with fitness – pleiotropy takes the form of a negative relationship called trade-off (Cody, 1966; Gadgil & Bossert, 1970; Stearns, 1989). Although trade-offs are usually detected by fitting simple regression models (*e.g.* Charnov & Ernest, 2006; Walker *et al.*, 2008; Mappes *et al.*, 2004; Roff *et al.*, 2002; Tucic *et al.*, 2005; Roff *et al.*, 2003; Andersson *et al.*, 2002; Hanski *et al.*, 2006), these relationships are not necessarily linear. Their precise shape (representing the standing genetic variation in a population) is actually a major evolutionary parameter that, by constraining movements on fitness surfaces, conditions evolutionary outcomes (Levins, 1968; de Mazancourt & Dieckmann, 2004; Leslie *et al.*, 2017; Verin *et al.*, 2017; Pisztor *et al.*, 2016). Trade-off shapes have long been considered inescapable constraints on the combination of traits that can exist, but this view is changing (Braendle *et al.*, 2011; Garland Jr & Carter, 1994). Indeed, recent empirical work has shown that trade-off shapes are highly context-dependent, responding plastically to experimentally manipulated environmental changes (Jessup & Bohannan, 2008; Maharjan *et al.*, 2013; Messina & Fry, 2003; Sgrò & Hoffmann, 2004). Moreover, trade-off shapes have been shown to change in a heritable manner, and evolve in organisms placed in experimentally manipulated environments (Roff *et al.*, 2002; Leroi *et al.*, 1994).

It is commonly accepted that negative relationships prevail because they result from the differential allocation of finite resources (Van Noordwijk & de Jong, 1986; Stearns, 1989; Agrawal *et al.*, 2010; Stearns, 1992; Roff, 1993; Harshman & Zera, 2007; Reznick *et al.*, 2000; Zera & Harshman, 2001). In multicellular organisms, resource allocation is regulated by hormones, whose joint effect on several traits creates so-called hormonal pleiotropy (Finch & Rose, 1995). For example, the internal concentration in juvenile hormone JH specifies the position – *i.e.* the combination of traits – along trade-offs between energetically-reliant traits like fecundity and survival in *Drosophila melanogaster* (Flatt, 2005) or fecundity and dispersal in *Gryllus firmus* (Zhao & Zera, 2002). Due to the key role of hormones as mediators of trade-offs, changes in the endocrine system are good candidates to explain the aforementioned plastic and heritable changes in trade-off shapes (Ketterson & Nolan Jr, 1992).

The question of how the endocrine system changes and responds to selection is addressed empirically by the emerging field of evolutionary endocrinology (Zera & Zhang, 1995; Cox *et al.*, 2016a,b; Garland *et al.*, 2016). This field, unfortunately, lacks a theoretical companion that would predict how endocrine systems should evolve depending, for instance, on the ecological context. The closest theory that has addressed related questions to our subject originated with van Noordwijk and de Jong’s seminal paper (Van Noordwijk & de Jong, 1986), and was later improved by dynamic energy budget models (Kooijman, 1986, 2000; Jager *et al.*, 2013). These models consider ‘mutations’ that modify resource allocation, thereby ignoring the actual regulatory and coding mutations impacting the proteins that form the endocrine system. Therefore, despite its momentous contribution to our understanding of evolution, this theory has presumably limited power to explain how the endocrine system and trade-off shapes should evolve.

Our aim with the present study is to initiate a reflection on how endocrine systems evolve and how this evolution impacts the trade-off shape. In this spirit, we built an evolutionary model where the interactions between hormones and receptors control the energy allocation to two trait-converting structures. Mutations in this model can change the conformation and expression of hormones and receptors. We show that, through the appearance and fixation of these mutations, the shape of the trade-off between the two traits does indeed evolve. Furthermore, this shape is highly dependent on a rarely considered parameter, the cost associated with resource storage. Consequently, in the simple ecological setting considered here, evolutionarily expected trade-off shapes and their underlying endocrine mechanism should depend both on the resource that is traded, and on parameters that set the burden associated with storage structures

## TRADE-OFF SHAPES DO EVOLVE

The conversion of an energetic resource into trait values is carried out by two specialized structures in our model, whose efficiency decreases as the inward flow of the resource increases (see figure 1). This flow is contingent on the dynamics of hormone-receptor binding at the surface of these structures, as newly formed hormone-receptor complexes activate inward transporters of the resource (see the *Material and methods* section for a precise, mathematical description of the model). Thus, following a meal – suddenly increasing the internal concentration of the resource – the resource may enter the converting structures, or a storage structure insofar as the internal concentration of the resource is above a threshold (figure 1). Past this threshold, the resource is instead released from the storage structure to maintain a constant internal concentration. Storage comes with a cost, such that the amount of the resource released may be lower than the amount that had actually been stored. After the storage structure has gone empty, the internal concentration of the resource decreases until it reaches a critically low value (figure 1). At this point, each individual’s fitness is calculated as the sum of its trait values.

**Figure 1:**
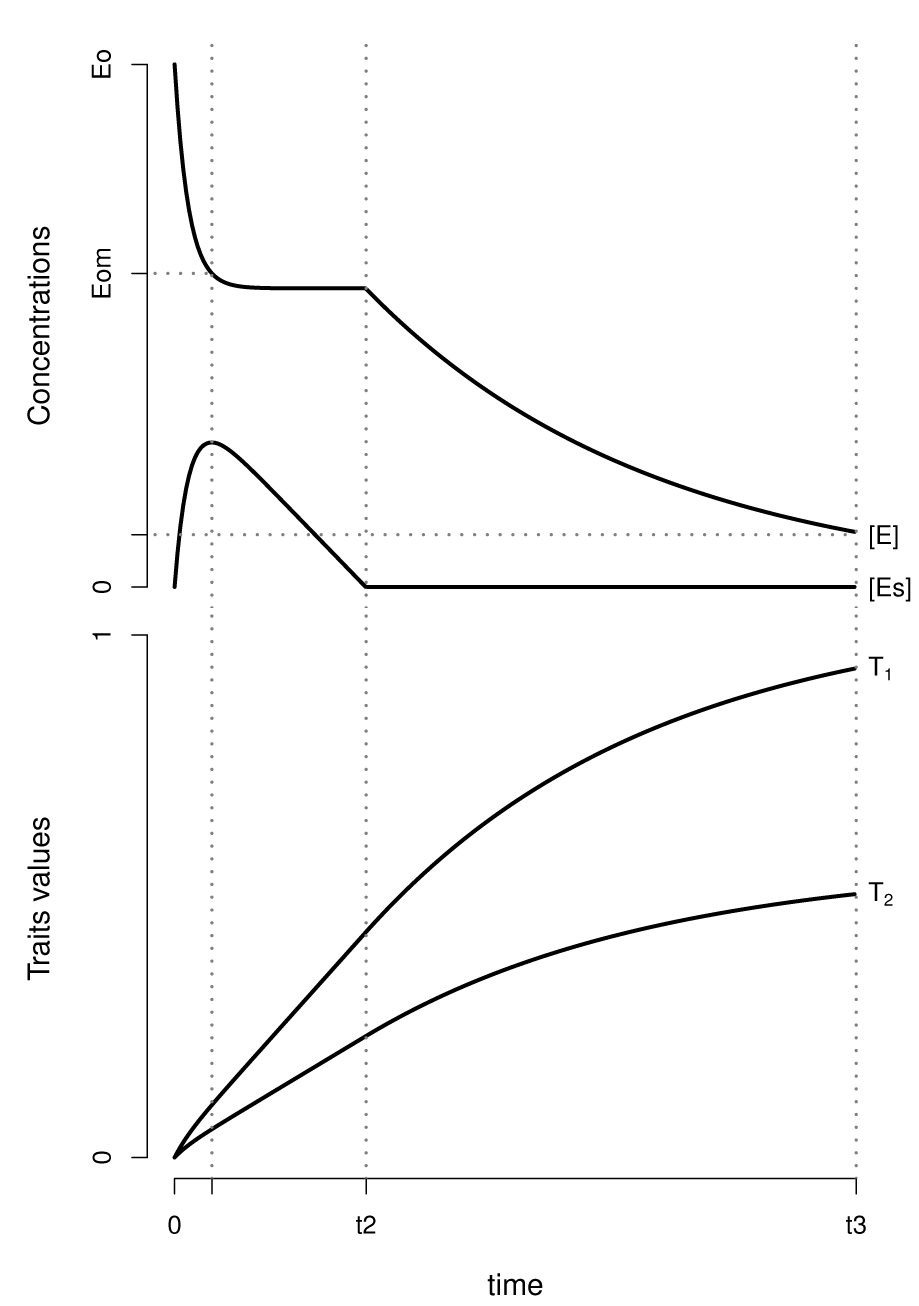
The final trait values, T_1_ and T_2_, depend on the dynamics of the concentration of the resource circulating the organism, [*E*], and stored, [*E*_*s*_]. [*E*] starts at Eo immediately after a unique meal, and decreases as the resource is both stored ([*E*_*s*_] increases) and used by the two converting structures (*T*_1_ and *T*_2_ increase). As [*E*] decreases below a threshold *E*_om_, the resource is released from the storage structure ([*E*_*s*_] decreases), which compensates for the energy used and converted into traits Ti and T_2_. When no stored resource remains ([*E*_*s*_] = 0), [*E*] inevitably decreases (as *T*_1_ and *T*_2_ keep increasing), until it reaches a minimum threshold. The organism’s final trait values *T*_1_ and *T*_2_ calculated at this point are used to assess its fitness. The hormones and receptors at the surface of the two converting structures control their resource intake, and thereby also the dynamics of [*E*] and [*E*_*s*_], such that mutations of the expression and conformation of these proteins may change the organism’s phenotype in a complex and unpredictable way. We used specific parameter values to obtain a clear illustration : *E*_o_ = 1, *E*_om_ = 0.6, *E*_min_ = 0.1, a = 0.0001, Cstorage = 0.1, Ω = 0.1, b = 0.002, ω = 0.1

When gene expression noise is ignored, an indivi-dual’s phenotype is entirely contingent on its genotype. Four independent genes code for two hormones and two receptors, whose expressions and conformations may change, respectively due to regulatory and coding mutations. These two types of mutation may impact the hormone-receptor dynamics, with potential effects on the resource allocation dynamics and thus on the trait values. These mutations change in frequency through the combined effect of drift and selection. We apply directional selection on each trait, a common regime among traits relying on resource allocation (e.g. ferti-lity or juvenile survival).

Our results show that trade-offs can, in principle, evolve. A representative example of the ongoing evolutionary dynamics is represented on figure 2-a. The population is initially monomorphic, so its occupies a single point on the phenotype space with low values of traits 1 and 2. As genetic variation builds up in the population, the phenotypes spread along an almost linear relationship (t = 100 in fig. 2-a). Around timestep 1300 trait 1 increases following a trajectory nearly parallel to the *x*-axis – in about half the simulations trait 2 instead increased. This change conveys a very sharp fitness increase because the population reaches a different isocline in the fitness landscape. Then the population roughly follows this isocline toward a point where trait 1 equals trait 2 (t = 5000 in fig. 2-a). This move in the phenotype space is actually associated with a slight increase in fitness, due to the higher efficiency of resource conversion when the two trait-converting structures are equally employed.

**Figure 2:**
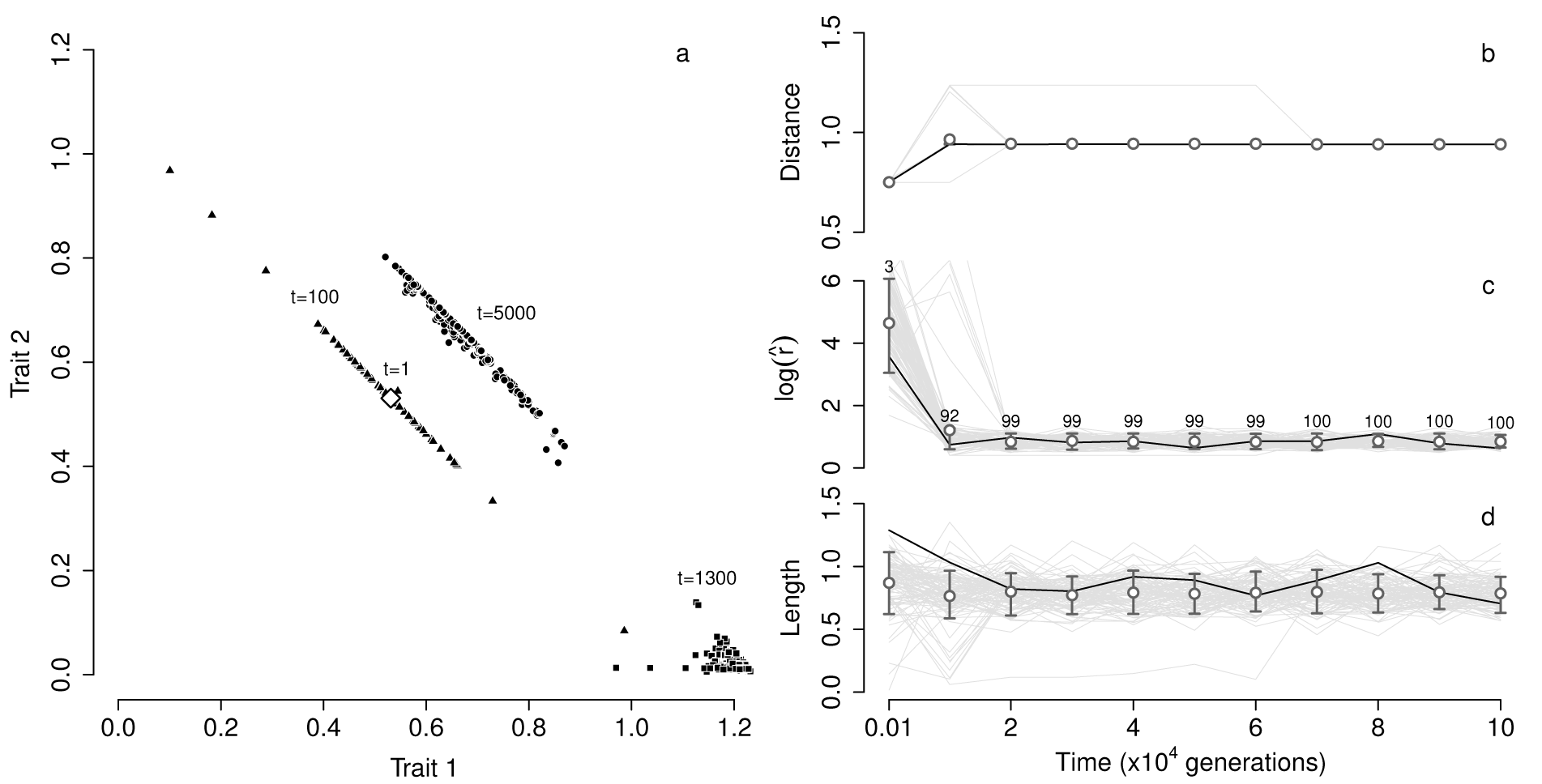
Trade-off shapes evolve in simulations where mutations change hormones’ and receptors’ conformation and expression, and selection acts depending on the resulting trait values. Panel a shows a few important timesteps in one representative simulation. We defined three parameters to study changes in shape systematically in an ensemble of 100 identically parameterized simulations, by fitting a circle to sets of points at regular timesteps (see text and *Material and methods section*). Panel b shows the temporal dynamics of the distance of the median projection on the fitted circle to the origin. The natural logarithm of the fitted circle’s radius (inversely related with the trade-off curvature) is represented in panel c, and the length of the trade-off (*i.e.* the distance along the circle between the two most distant projections) is represented in panel d. 99 replicate simulations are represented in grey in panels b-d, and the simulation in panel a is outlined in black. Dots and bar represent respectively the mean and quantiles (0.1 and 0.9) of each parameter’s distribution (across simulations) at each timestep. Standard parameter values where used (as defined in the *Material and methods* section), and *C*_storage_ = 0.5.

We identify changes in the shape of the trade-off by fitting a circle to the combinations of traits that coexist at any time point in our simulations (see the *Material and methods* section and figure 3). We projected each individual datapoint on the fitted circle, and calculated the Euclidean distance *d* of the median projection to the origin. Because d is closely related with fitness, it increases during the simulation (figure 2-b). The trade-off curvature decreases as the circle’s radius 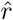 increases; we thus use log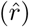 as a negative proxy for curvature. Individuals in the population are initially distributed along a fitness isocline, such that the trade-off is almost linear (as shown by the fitted circle’s high radius ; fig. 2-c). Then the curvature rapidly increases in all the simulations, toward 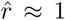. At the end of the simulation, all estimated trade-offs are concave (the median projection is closer to the origin than the center of the line between the two most extreme projections). We also estimate the trade-off length as the distance between the two most distant projections on the circle, whose between-simulation mean quickly stabilizes around 0.6 (figure 2-d).

**Figure 3:**
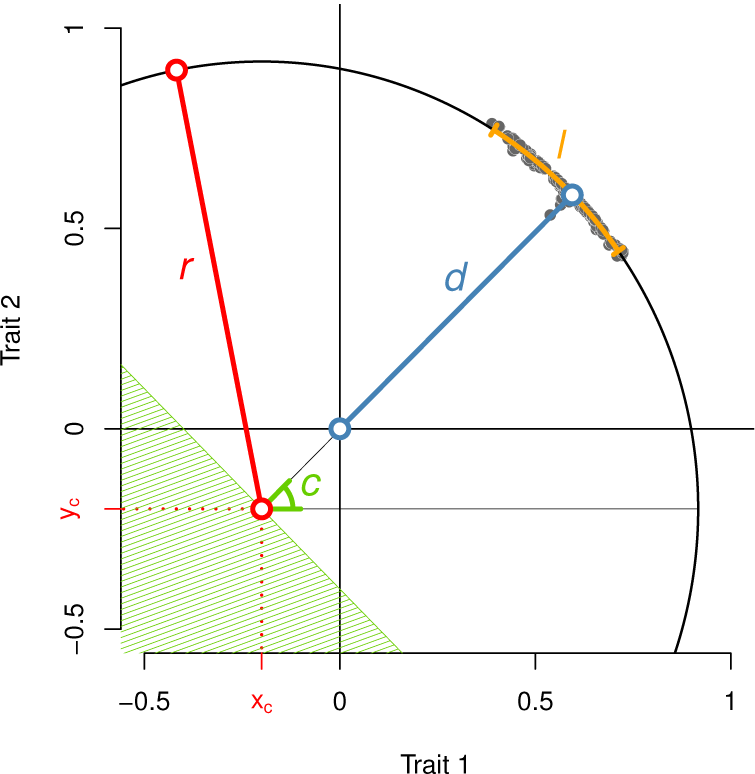
We extracted four descriptive parameters by fitting a circle to the phenotypes (trait 1, trait 2) at the end of our simulations. The parameters are l (the length of the trade-off – in orange), *d* (the distance to the origin-in blue), 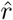 (the circle’s radius – in red) and *c* (the median angle, from which we determine the concavity of the trade-off – in green). We minimized the squared distances of the orthogonal projection of individual datapoints to obtain 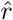 and the center of the circle (*x*_*c*_, *y*_*c*_). Then we calculated the angle *c* as the median among the angles calculated for all datapoints. The trade-off is considered convex if *c* is between 3 π /4 and 7 π /4 (green area) and concave otherwise. We calculated *d* as the distance between the point on the fitted circle with angle *c* and the origin. *l* is the distance along the circle between the two points with the most extreme angles (see the *Material and methods – Estimates of shape parameters* section for a precise method description)

We ran neutral simulations, where individuals have equal fitnesses regardless of their phenotype, with identical parameters as in figure 2 (figure S1). The difference in evolutionary outcomes is striking : in the neutral case, all trade-offs are convex (while all are concave under selection), and trade-offs are more linear and longer on average.

## THE EVOLUTION OF TRADE-OFF SHAPES DEPENDS ON STORAGE COST

As mentioned above, our model includes a parameter C_storage_ for the cost of storage, which in nature comes from two distinct sources. First, large molecules need to be produced for storage and later broken down at the time of release (Hers, 1976; Meyers, 1995; Alonso *et al.*, 1995); second, maintaining storage structures can be costly and the added weight and volume might have negative consequences for the organism’s fitness. The cost of producing storage molecules is likely dependent on the resource : various types of molecules are ingested during a meal, with specific storage constraints (Williams *et al.*, 1963). Carbohydrates like glucose, for instance, are stored under the form of glycogen, which is rapidly converted back into an energetic resource but requires a large amount of water – and thus a large volume and weight – for storage, contrary to lipids that are stored as fatty acids (Schmidt-Nielson, 1997). Specific advantages and disadvantages associated with each resource may translate into specific storage costs, likely in interaction with the ecology of the species considered. For example, the added weight might have a moderate fitness impact for marine species compared to aerial ones.

We varied this cost between 0 and 1 and found that the evolutionarily expected shape parameters drastically change in response (figure 4). As the cost increases, populations reach lower trait values (and are therefore closer to the origin, fig. 4-a), as an increasing part of their resources are inevitably wasted. Presumably, the genotypes that evolve under a specific storage cost optimize the speed of resource consumption. Faster consumption would lead to a less efficient energy conversion, while slower consumption would require storing more resources, and paying the associated cost. Under a high storage cost, selection thus favors rapid converters (as shown in fig. S2) that allocate a similar amount of energy to each trait. A deviation from this ideal allocation decreases the efficiency of the most-used converting structure, which explain that the trade-off becomes more curved – *i.e.* that log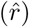 decreases – as the storage cost increases (fig. 4-b). The trade-off length, representative of the population’s standing genetic variation, decreases conjointly (fig. 4), likely because extreme individuals on these highly curved trade-offs have low fitnesses.

**Figure 4:**
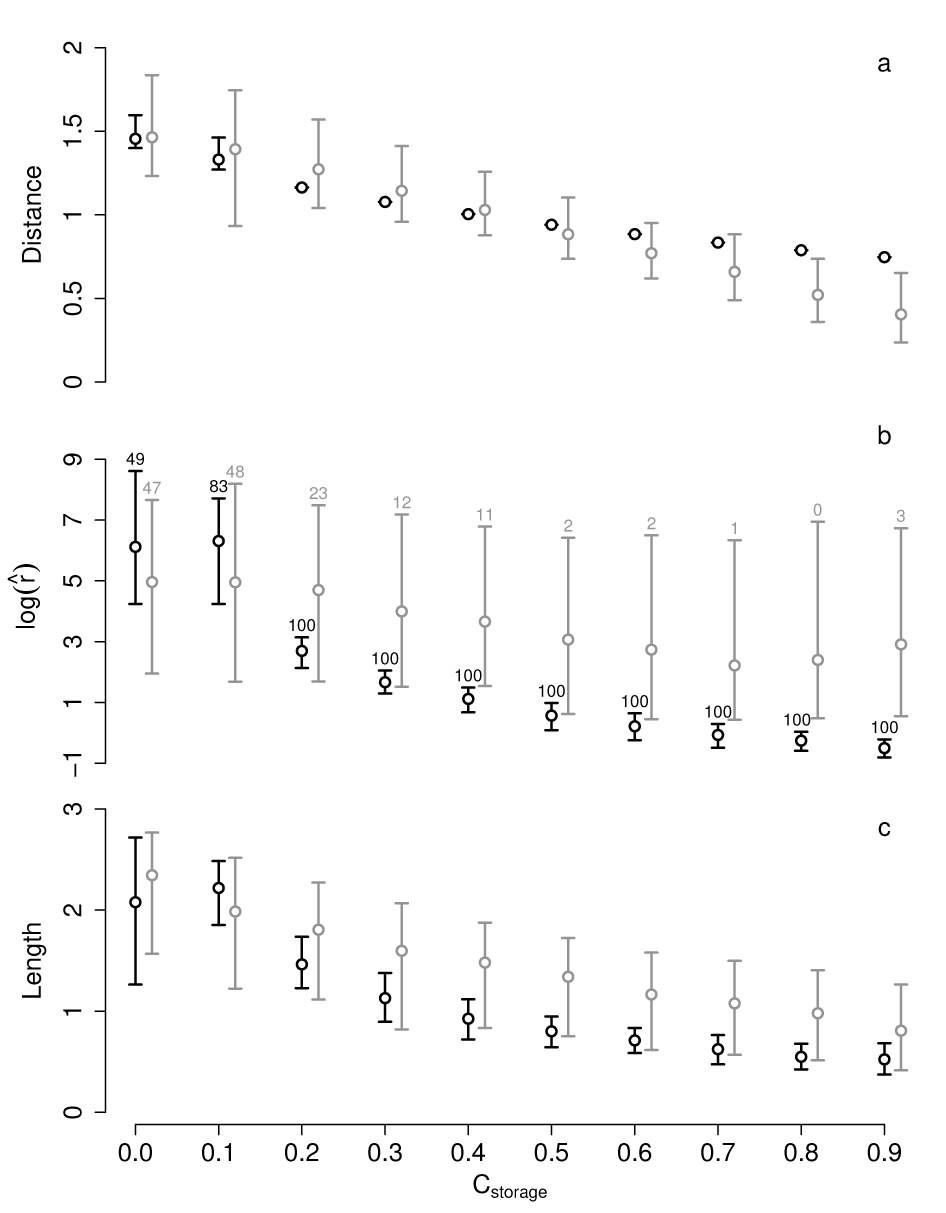
Evolutionarily expected shape parameters depend on the cost associated with storage *C*_storage_, due to selection. The impact of selection appears by comparing the evolutionary expectations of a model with directional selection on both traits (in blue) with those of a neutral model (in grey; individuals have identical fitnesses in this model, independent of their trait values). Definitions for the shape parameters can be found in the text, in the *Material and methods* section, and in figs. 2,3.

A comparison of these results with those of a neutral model indicates that selection plays a major role in the evolution of trade-off shapes. Surprisingly, at low storage costs the neutrally expected distance *d* is higher on average than under selection (fig. 4-a). This is likely due to the absence of selection for an optimum conversion speed, which maintains populations on the line where trait 1 equals trait 2. For similar reasons, the curvature of trade-offs remains independent of the storage cost (fig. 4-b) in the neutral case, leading to marked differences with the results under selection at high storage costs. The trade-off length decreases slightly as the storage cost increases in the neutral model (fig. 4-c) – possibly because individuals are closer to the origin in this situation, limiting the range of positive trait values along the trade-off.Trait 1

We have performed additional sensitivity analyses to three parameters : the population size, the mutation rate and the level of gene expression noise (figs S3-S5). All three parameters increase the variation among individuals and thus the length of the trade-off, and in the case of noise this increase in length coincides with a slight decrease in the curvature of the trade-off, which may have two distinct causes. First, more linear trade-offs may be selected as they make the organism robust to noise (Wagner, 2005; Draghi & Whitlock, 2015). Indeed, if the trade-off is more linear and aligns with the fitness isoclines, perturbations of gene expression levels still move individuals along trade-offs, but these moves have moderate fitness consequences. Second, noise has the effect of diminishing the strength of selection (Mineta *et al.*, 2015). Hence, as noise increases in our simulations, it is possible that selection on the curvature weakens, such that the curvature decreases towards the neutral expectation (see figure 2). Although we cannot entirely rule out this second hypothesis, the reduced variance among simulations under strong noise suggests that robustness may play some role in flattening trade-offs.

## THE LEVEL AT WHICH EVOLUTION CONVERGES

At this point we need to introduce the classical heuristic of genotype networks, which are made of nodes (each one a genotype) connected by mutations or recombination events (fig. 5; Smith, 1970; Draghi *et al.*, 2010; Rajon & Masel, 2013). Typically, a genotype has a restricted number of neighbors relative to the total number of nodes in the network. A population with standing genetic variation (SGV) occupies a certain number of nodes on the network. Since SGV arises from short-lived mutations and recombination events (Barrett & Schluter, 2008; Lerouzic & Carlborg, 2008), the population likely occupies a small subset of the whole genotype space at any point in time. This subset defines the features of SGV, including a trade-off’s shape. In the longer term, a population might make larger moves in the genotype space, changing its mutational neighborhood and thereby, possibly, the shape of its trade-off(s).

**Figure 5:**
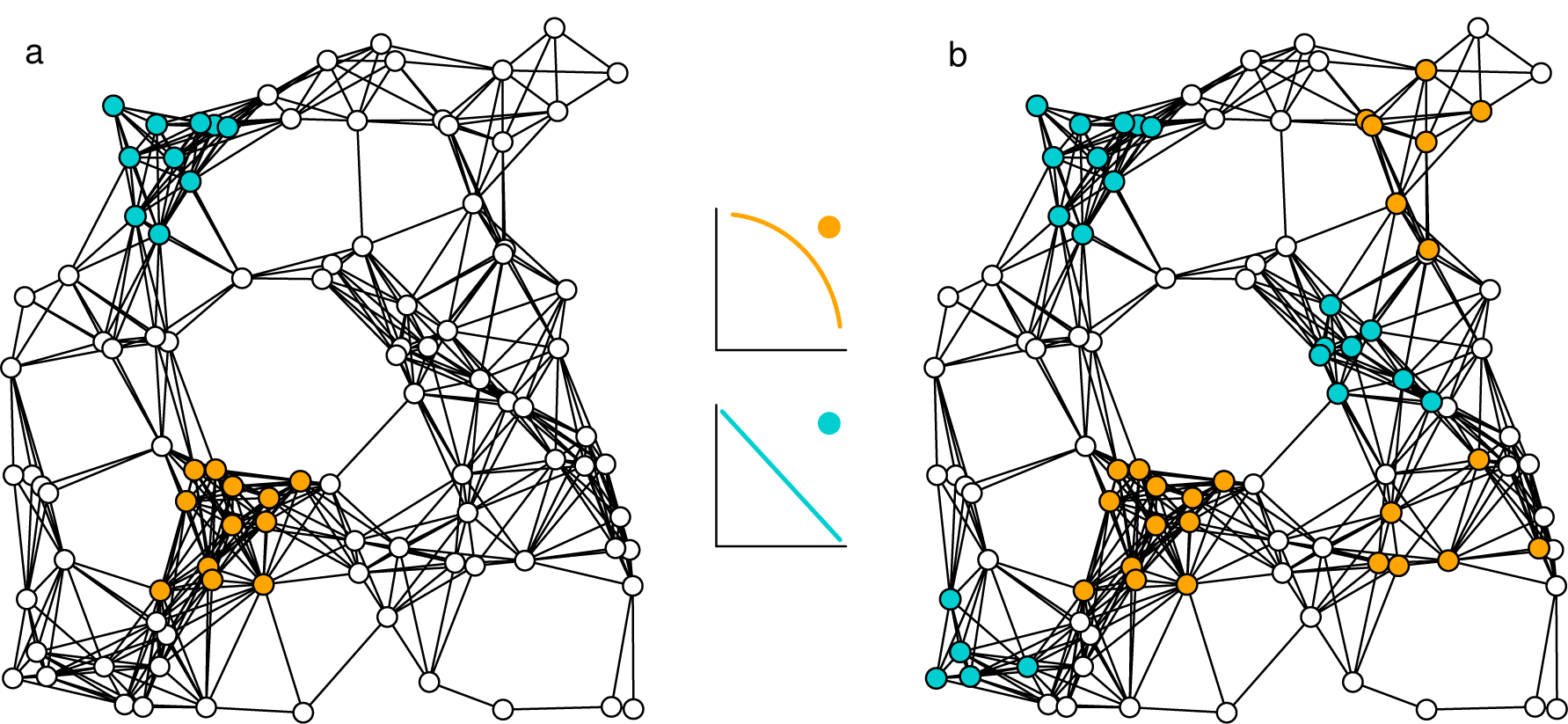
Two genotype spaces illustrate the two hypothesized relationships between the genotype (and hence the physiological mechanism) and the phenotype. Genotypes are represented by nodes, linked by mutations or recombination events. The phenotype is specified by a color for some genotypes, corresponding to the trade-off shapes represented in the center of the figure. In panel a, the two colored mutational neighborhoods are unique solutions to generate a given trade-off shape (no redundancy), whereas in panel b several mutational neighborhoods can lead to similar phenotypes (redundancy at the genotype level).

As we have seen, *e.g.* in figure 2, evolution is clearly convergent at the phenotype level – similar trade-off shapes evolve in similar contexts. We next ask whether evolution is also convergent at the genetic and physiological levels. As anticipated by Garland and Carter (Garland Jr & Carter, 1994), convergence at these low levels would mean that organisms confronted to similar problems evolve identical mechanisms. This means that a unique neighborhood on the genotype space should yield a particular trade-off shape, as represented in figure 5-a.

In order to test this hypothesis, we used multivariate analyses on two sets of variables extracted from our simulation endpoints : a genotype set, with the expressions and conformations of hormones and receptors ; and a physiological set that groups different genotypes yielding similar physiological mechanisms – for instance, we would group a genotype whose hormones have conformation 25 and receptors 35 with one with conformations 5 and 15, respectively, all else being equal. see details in supplementary text 1). We first performed a Principal Component Analysis (PCA), which revealed no apparent association between the variation in the genotypic or physiological variables and the storage cost (fig. S6). It indicates that the variation between populations in a similar context (*i.e.* with the same *C*_*storage*_) is large compared to the variation between contexts. This means that evolution has found a variety of genetic and physiological solutions when confronted to a similar problem.

Despite the general lack of convergence at the genetic and physiological levels, it is possible that the physiological systems that evolve in a particular context have some common characteristics. This hypothesis can be tested by performing a Canonical Correlations analysis (CCA), aimed at identifying the combination of variables most correlated with the storage cost. We found a significant correlation between the storage cost and a linear combination of the physiological dataset (Wilk’s A = 0.93, *p* = 5,09.10^−12^), which as expected explains a small part of the overall variance (*r*<sup>2</sup> = 0.0676). Studying the contributions of the variables to this linear combination shows that as the storage cost increases, hormones and receptors have higher expression rates but, surprisingly, not better affinities on average (Text S1 and fig. S7).

These results lead to the conclusion that evolution is not convergent at the genotype level, and scarcely so at the physiological level. Back to our genotype space heuristics, this means that several subsets of this space produce similar phenotypes, as illustrated in figure 5-b. At first sight, this prediction seems incompatible with empirical observations indicating that major regulators in endocrine mechanisms are strongly conserved (Tatar *et al.*, 2003; Barbieri *et al.*, 2003), suggesting that evolution has found few solutions to a given problem. However, this observation could also be explained by a specific structure of the genotype space preventing large evolutionary moves between similar neighborhoods. This situation can be pictured in the hypothetical genotype space in figure 5-b by giving genotypes represented by white dots very low fitnesses : changes in trade-off shapes are still possible (some ‘orange’ genotypes have ‘blue’ neighbors) but reaching distant neighborhoods is highly unlikely.

## CONCLUSION

In our model, mutations have cascading effects on phenotypic traits through changes in the physiological systems. The main benefit of this approach is that it make movements on the genotype space (either through mutation or recombination) and the effects of environmental noise more realistic. It prevents the modeler from making arbitrary assumptions about the distribution of mutational effects, as is common in evolutionary ecology models. The debate over the introduction of mechanistic assumptions into evolutionary models dates from the origins of the modern synthesis, when Wright argued against Fisher in favor of physiologically realistic models to explain the prevalence of recessive mutations (Fisher, 1928; Wright, 1929, 1934). Despite today’s general acceptance of a major role of physiology in the evolution of dominance (Bourguet, 1999), Wright’s proposal of introducing physiological details in evolutionary models has remained largely ignored.

Not only are physiologically-driven evolutionary models more realistic to some extent, they also allow to ask novel questions, such as ‘how does the endocrine system evolve?’ or ‘why is it so complex?’. The question of the evolution of a trade-off’s shape that motivated this study could not be answered properly without this approach, since this shape necessarily emerge from the underlying physiological system. We have found that selection plays a crucial role in this evolution : neutrally, a population would move on the genotype space towards locations where the trade-off is generally convex and flat, while selection makes the trade-off concave and curved. Importantly, the impact of selection depends on context, such that the evolutionarily expected trade-off shape depends on characteristics of the resource that is traded and on the ecology of the species.

## MODEL

The model considers the conversion of an energetic resource by specialized structures into two traits under directional selection. Hormones may bind a receptor on a converting structure, which activates inward transporters of the resource. We assume that the resource intake by the organism (“meal” thereafter) takes place once the hormone-receptor dynamics has reached its unique equilibrium (studied in subsection). The model then considers the dynamical absorption of the resource by the two converting structures as well as exchanges with a storage structure (subsection). The energy accumulated by structures is immediately converted into two traits values (subsection). We let this physiological mechanism change by mutation and evolve under the influence of selection and drift as described in subsection, and analyze these simulation results as described in subsection.

### Hormone-receptor binding dynamics

Hormones are produced through the expression of two genes (identified by the subset *k ∈* {1, 2}) by a specialized structure, followed by the export of their products. Individuals are diploid, so two different hormones can be expressed by each gene (*p ∈* {1,2}). The internal concentration of the hormone expressed by the allele *p* of gene *k* is denoted [H*_kp_*]. Another structure degrades the hormones, depending on their concentration. Two structures converting energy into traits express receptors on their surface. The concentration of the receptor expressed by the allele *q* of gene *i* (with *q* and *i* ∈ {1,2}) on structure j (with *j ∈* {1, 2}) is denoted [*R*_*iqj*_] (we ignore the specific processes of production and degradation of receptors). Hormone kp and receptor iq form complexes at rate 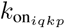 (depending on their respective conformations, see *Hormone-receptor binding affinity*), and complexes dissociate at rate k_off_. The concentration of complexes formed between receptor iq and hormone kp on structure j is denoted [*R*_*iqj*_*H*_*kp*_]. The hormone-receptor binding dynamics are described by the generic equations [1]-[5], where *n*_*h*_, *n*_*r*_, *n*_*s*_ and *n*_*ch*_ are respectively the numbers of hormone- and receptor-coding genes, converting structures, and chromosomes. We set *n*_*h*_ = *n*_r_ = *n*_*s*_ = 2, such that one possible evolutionary outcome is that the energy allocation to each converting structures is regulated by one specific couple of hormone and receptor.

#### Free circulating hormones

The concentration of hor-mone *kp*, [H*_kp_*], increases as the hormone is produced at a rate *α*_*kp*_ (first term in equation (1)) and as hormone-receptor complexes dissociate at a rate koff. All n_r_ receptors on each structure are considered in the second term of equation [1]. Conversely, [*H*_*kp*_] decrease as complexes form at rate 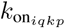-specific of the receptor expressed by the allele *q* of gene *i* and of the hormone expressed by the allele *p* of the gene *k* – and due to unspecific hormone degradation (last term).

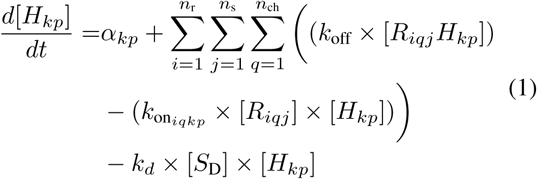

#### Free receptors

The number of free receptors increases as complexes dissociate and decreases as complexes form with each of the *n*_*h*_ hormones produced by the organism (equation (2)).

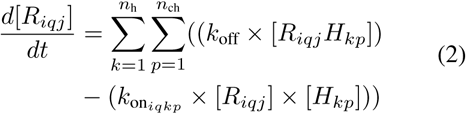

#### Hormone-receptor complexes

The number of hormone-receptor complexes increases as hormones bind on receptors and decreases as complexes dissociate (equation (3))

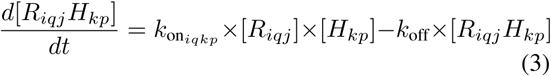

The concentration of receptor iq is constant 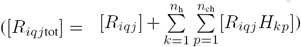, such that each dissociated complex releases a free receptor and each formed complex monopolizes a free receptor.

#### Degradation structure

Our model includes a structure that degrades hormones non specifically through endocytosis. The liver can be considered an example of such structure, which degrades the growth hormone (Baumann *et al.*, 1987), insulin (Duckworth *et al.*, 1998) and glucagon (Deacon, 2005). The dynamics of the concentration of free degradation sites [*S*_D_] and of degradation sites occupied by hormone *kp*, [*S*_D_H*_kp_*] are described by equations [4] and [5]. The first term in each of these equations relies on the assumption that degradation sites are instantly freed.

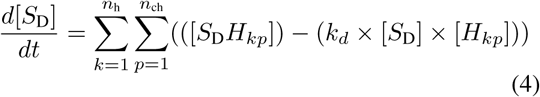

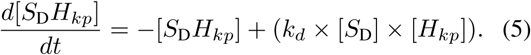

We set the total number of degradation sites 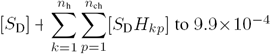 to 9.9 x 10^−4^, and the rate of hormone internalization *k*_*d*_ to 4.10^−5^ (Pearce *et al.*, 1999); we did not allow these parameters to change by mutation. other parameters in the model, however, can mutate and change the equilibrium concentration of the free hormone (e.g. the rate of production of each hormone). As described in the section *Evolutionary and population dynamics*, we randomly drew the values of mutable parameters at the beginning of each simulation from known distributions, and verified that the internal hormonal concentration at equilibrium is, initially, within the range 5.10^−12^ — 10^−8^ M, in accordance with experimental data (Polonsky *et al.*, 1988; Zadik *et al.*, 1985).

#### Hormone-receptor binding equilibrium

Hormone-receptor binding dynamics eventually reach an equilibrium where [*H*_*kp*_], [*R*_*iqj*_] and [*R*_*iqj*_ *H*_*kp*_] are stable (equilibrium values are identified by a star in equations (6)-(8), obtained by solving the system formed by equations (1)-(5) (see details in supplement).

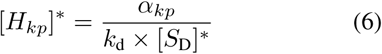

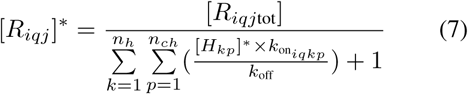

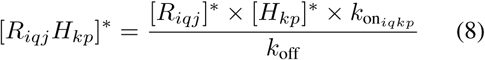

Despite some simplifying assumptions detailed below, we consider that receptors in our model function like G protein coupled receptors, whose activation produces an internal signal for a limited amount of time, followed by their internalization and recycling into free receptors. Hormone-receptor complexes normally dissociate at a rate k_off_ of the order of 10^−4^ — 10^−5^ (Pearce *et al.*, 1999). We voluntarily over-estimated this rate (*k*_off_ = 0.1) to model recycling, thereby considering that both the hormone and the receptor are released when recycling is completed. With this parameterization, a receptor is recycled within 10 minutes on average, which is biologically realistic (Von Zastrow & Kobilka, 1992).

Considering that both the hormone and the receptor are released is mathematically convenient but unrealistic, as recycling is most often followed by the degradation of the hormone. We verified that this assumption does not impact our results (see SI figure S8) using numerical simulations where only the receptor is recycled – *i.e.* the hormone is not released.

### Energy allocation dynamics

An individual takes a unique meal at t_0_ which sets its internal concentration of an energetic resource [*E*] to *Eo* = 1. Then [*E*] decreases, as the resource is partly stored and partly allocated to the two structures converting it into traits values. In our model, newly formed hormone-receptor complexes instantaneously stimulate transporters of the resource and become inactive. The number of complexes formed on structure *j* per time unit at equilibrium equals :

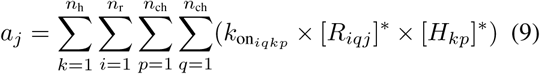

which we multiplied by a constant *C*_f_ = 1000 to obtain the inward flow of energy for this structure.

The energy allocation dynamics consider temporal changes in [*E*] as well as in [*E*_*s*_], the concentration of the resource stored. The ODEs for the instant changes in [*E*] and [*E*_*s*_] are given below (equations [10]-[11]) :

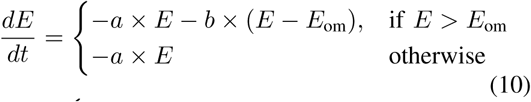

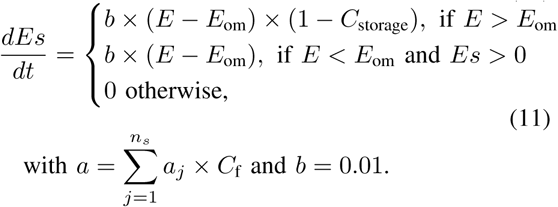

with 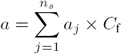 and *b* = 0.01.

We solved these equations (see SI text 2) and obtained the temporal dynamics, [*E*](*t*) and [*E*_*s*_](*t*) by separating them into three phases illustrated in figure 1. Phase 1 starts at t_0_ (the meal) and goes on as long as [*E*] > *E*_om_, *E*_om_ = 0.08 being a concentration threshold above which energy is stored. During phase 2 – which starts at ti (when [*E*] = *E*_om_) – the resource is released from the storage structure which compensates for its consumption by converting structures. Phase 3 begins when [*E*_*s*_] reaches 0, such that [*E*] decreases until it reaches the critically low value *E*_min_ = 0. 01.

### Energy conversion into traits

The energy allocated to each structure is converted into an increase of trait value 1 (for structure 1) and trait value 2 (for structure 2). Traits are deliberately abstract, but their biological relevance is quite obvious. For example, energy allocation to reproductive structures can increase the quality and number of gametes, thus increasing fertility. Likewise, energy allocation to somatic maintenance or camouflage may increase survival. Here we assume that allocating a lot of energy to one structure per time unit decreases the efficiency of conversion. The observation that fast feeding larvae of *D. melanogaster* have lower fitness than slow feeding larvae may support this assumption (Foley & Luckinbill, 2001; Prasad & Joshi, 2003; Mueller *et al.*, 2005; Roff & Fairbain, 2007), although it could also be partly explained by increased efficiency of digestion in slow feeding larvae. It is obvious, nonetheless, that some loss will occur as energy expenditure increases, because no organism is capable of instantly using large amounts of resources. We model this loss as :

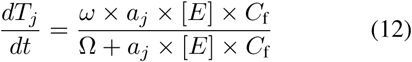

*Tj* is the trait value for structure *j*, ω = 0.02 is the maximum increase of trait value per *dt*, and Ω = 0.01 sets the speed with which this function saturates. We found the analytical expression of *T*_*j*_ (*t*), which follows the dynamics of [*E*](*t*) described above (see figure 1). The combination of trait values calculated at time *t*_3_ is the phenotype of the individual, used to calculate its fitness (see below).

### Evolutionary and population dynamics

The population is formed of 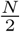 females and 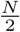 males. Offspring production is modeled using a Wright-Fisher process (Fisher, 1930; Wright, 1931), meaning that the population is entirely renewed at each timestep and that each newborn’s mother and father are sampled from the female and male pools, respectively. Each time an individual is chosen as a parent, the gamete it transmits to its offspring is formed by sampling independently one allele for each of the four genes (two hormones and two receptors) from the two parental chromosomes. The female status in the next generation is attributed to exactly *N/*2 randomly sampled individuals; the other *N/*2 become males.

In the neutral instance of our model, the probability that mother or father *i* is drawn equals 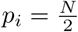In the model with selection, the probability *p*_*i*_ that parent *i* is drawn is proportional to its fitness *W*_*i*_ relative to other same-gender individuals in the population :

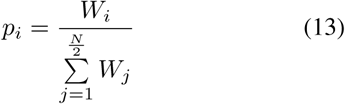

where *W*_*i*_ = *T*_1i_ *+ T*_*2i*_. Therefore, this model includes genetic drift (a genotype can go extinct by chance) and selection.

As stated above, the genotype in our model comprises four independent genes encoding two receptors and two hormones. Allele *q* at the gene coding for receptor *i* determines its expression on each structure *j*, [*R*_*jgj*tot_], and its conformation Ψ*R*_*iq*_. Similarly, allele *p* at the gene coding for hormone *k* sets its production rate *α*_*kp*_ and its conformation Ψ*H*_*kp*_. [*R*_*iqj*tot_] and *α*_*kp*_ directly affect the hormone-receptor dynamics in equations (1)-(8), and thereby the energy allocation dynamics. The receptor and hormone conformations (*ΨR_iq_ and ΨH_kp_*) are modeled as positive integers in our model. They also impact the aforementioned dynamics through their contribution to the specific on-rate constant, 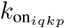 defined by equation (14) :

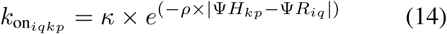

The affinity between hormone kp and receptor iq is maximum when Ψ*H*_*kp*_ = Ψ*R*_*iq*_, and decreases as the difference between these numbers increases.

#### Mutations

An allele’s contribution to the protein’s expression or conformation can mutate with probability μ = 10^−4^ per regulatory/coding sequence per generation. Mutations can change a receptor’s concentration [*R*_*ij*_tot] or an hormone’s production α_k_. In this case, the mutant’s value is multiplied by 10^γ^, where γ is sampled from a normal distribution with mean 0 and standard deviation σ*_m_* = 0.5. Coding mutations can change the conformation of an hormone or a receptor by adding an integer sampled from a discretized normal distribution with mean 0 and standard deviation 1, and from which 0 values were excluded to avoid neutral mutations. Conformations were, kept within the range {1; 1000} by considering the extreme values, 1 and 1000, as immediate neighbors.

#### Initial conditions

Each population at t = 0 is monomorphic, such that all individuals share the same genotype and are homozygotes for all genes. We nonetheless introduced variation in the initial genotypes across replicate simulations, by mutating every coding and regulatory sequence once from an average starting genotype. This average genotype produces hormones at rate α*_kp_* = 9.9 x 10^−10^ (*k,p* G {1,2}) and expresses [*R*_*iqj-tot*_] = 9.9 x 10^−5^ receptors of type iq at the surface of structure *j* (*i, q, j* G {1,2}), and has hormones and receptors conformations Ψ*H*_*1p*_ = 9, Ψ*H*_*2p*_ = 109, Ψ*R*_*1q*_ = 29, and Ψ*R*_*2q*_ = 129 (*p,q* G {1, 2}).

### Estimates of shape parameters

As described in the previous section, the physiological mechanism can change by mutations and evolve. We expect at least some of these mutation to have consequences on the shape of the trade-off between traits 1 and 2. We estimated these changes by fitting a circle to individual coordinates in the phenotype space formed by these traits. Because no causal relationship should be expected between trait 1 and trait 2, we performed an orthogonal regression to obtain estimates of the radius of the most fit circle 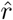 and the coordinates of its center (*x*_*c*_, *y*_*c*_) (Coope, 1993). From these estimates, we calculated three important parameters that quantify the curvature, the length and the position of the trade-off.

The curvature increases as the radius or its loga-rithm, 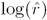, decreases – obtaining 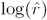 is straight-forward from the analysis. The length l – which is representative of the standing genetic variation in the population – is the distance between the two most distant orthogonal projections on the fitted circle. To obtain it, we first calculated the angle of each individual *i* as 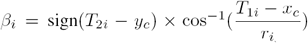, with *r*_*i*_ the Euclidean distance between (*T*_*1i*_, *T*_*2i*_) and (*x*_*c*_, *y*_*c*_) using a gradient descent method. Then we calculated 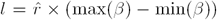. The position is evaluated by the distance of the median projection to the origin, d – a proxy for fitness, which we use in the neutral and in the selection model alike. We extracted the median angle in the distribution of β*_i_* β*_M_* and calculated the distance to the origin 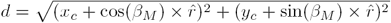.

